# Sensory processing reallocation from external to internal signals in REM sleep

**DOI:** 10.64898/2026.03.16.712081

**Authors:** Jacinthe Cataldi, Andria Pelentritou, Sophie Schwartz, Marzia De Lucia

## Abstract

The brain continuously integrates information from the external environment (exteroception) and the internal bodily milieu (interoception). How the balance between these two processing streams shifts across vigilance states with differing levels of environmental responsiveness, however, remains poorly understood. Here, we examined neural responses to external auditory and internal cardiac signals across wakefulness and REM sleep microstates - tonic and phasic REM - which are characterized by progressively reduced responsiveness to external stimulation. High-density EEG was recorded in healthy participants (n=25). Auditory evoked potentials (AEPs) and heartbeat evoked potentials (HEPs) served as indices of exteroception and interoception, respectively, and were compared across vigilance states. AEPs progressively decreased from wakefulness to tonic REM and were most attenuated during phasic REM. In contrast, HEPs were preserved across REM microstates and were enhanced relative to wakefulness, indicating sustained - and even amplified - processing of cardiac signals during REM sleep. To quantify the relative weighting of external and internal signals, we introduce an exteroceptive–interoceptive index, defined as the ratio of auditory to cardiac neural responses. This index decreased systematically across vigilance states, revealing a graded shift from externally oriented processing during wakefulness to internally oriented processing during phasic REM, with tonic REM occupying an intermediate position. Together, these findings demonstrate that while responsiveness to external stimuli diminishes during phasic REM, the brain continues to prioritize physiologically relevant internal signals. The exteroceptive-interoceptive balance may thus provide a novel, mechanistically grounded marker of altered consciousness, particularly informative in contexts where behavioural responsiveness cannot be assessed.

**Graphical abstract:** 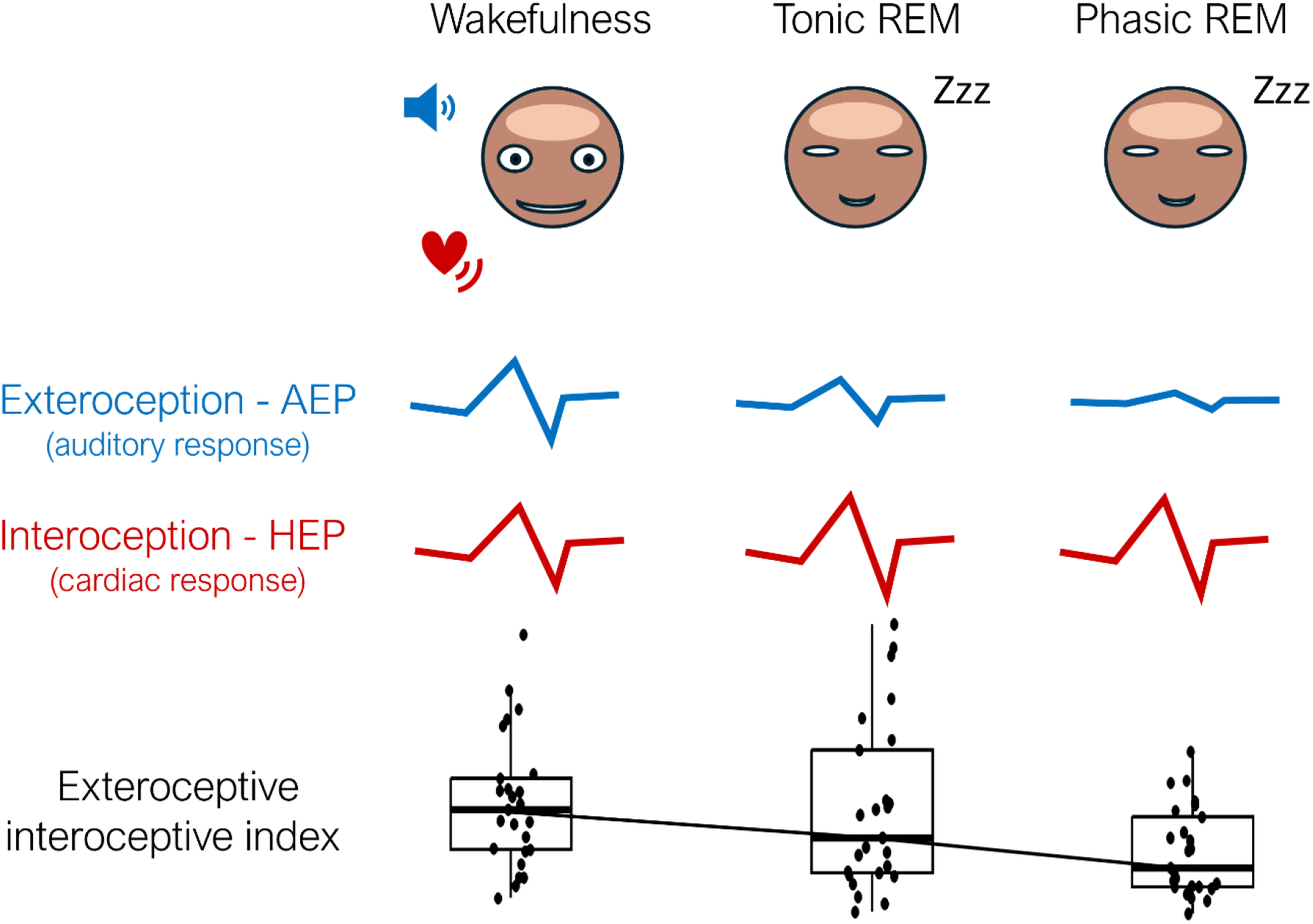

## Introduction

The brain continuously integrates sensory signals arising from the external environment (exteroception), and from within the body via interoceptive pathways^1-4^. Converging evidence indicates that these streams of information are not processed in isolation but are integrated to support functions ranging from physiological homeostasis to predictive inference about both external events and internal bodily states^5-8^. How the brain dynamically balances and prioritizes exteroceptive and interoceptive information remains, to date, unresolved.

A key aspect of this balance concerns the regulation of responsiveness - that is, how the brain detects, processes, and potentially reacts to external versus internal signals across different states of consciousness^9-11^. Wakefulness and rapid eye movement (REM) sleep provide an ideal framework for investigating this question. Although REM sleep exhibits electrophysiological features resembling wakefulness, combined with vivid internally generated experiences (i.e., dreams), it is also characterized by a marked disconnection from the external environment. Compared to wakefulness, REM sleep is associated with higher arousal thresholds and the absence of overt behavioural responses to external stimulation^12-14^.

Importantly, REM sleep is not a homogeneous state but alternates between two distinct microstates— tonic and phasic REM—which differ in physiological and autonomic features^15^. Tonic REM is characterized by moderate muscle atonia and relative physiological stability. Conversely, phasic REM features bursts of rapid eye movements, complete muscle atonia, transient muscle twitches, and autonomic variability^16^, with increased heart rate variability, irregular respiration, and greater sympathetic activation relative to tonic REM^17-19^. These microstates also differ in the degree of sensory disconnection: arousal thresholds to external stimuli increase and behavioural responsiveness decreases progressively from wakefulness to tonic REM and further to phasic REM^15,20^.

The progressive reduction in external responsiveness raises a fundamental question: does diminished responsiveness to external stimulation reflect a global reduction in sensory processing capacity, regardless of whether sensory inputs originate from the environment or from the body? If so, both exteroceptive and interoceptive processing should be attenuated during REM sleep, with the strongest suppression during phasic REM^18,21^. Alternatively, reduced responsiveness to external inputs may entail a flexible reallocation of processing resources, whereby the neural processing of internal bodily signals is preserved - or even enhanced - during states characterized by sensory disconnection from the environment.

Here, we tested these competing possibilities by investigating the neural responses to both external and internal stimuli in REM sleep, differently from previous work where these responses were studied in isolation^22-29^. External sensory processing was indexed by auditory evoked potentials (AEPs), whereas interoceptive processing was indexed by heartbeat evoked potentials (HEPs). Based on previous work, we predicted that AEPs would be attenuated during phasic REM relative to tonic REM and wakefulness^22,23,25,27,28^. In contrast, we hypothesized that HEPs would be enhanced during REM sleep compared to wakefulness, reflecting a potential shift of neural resources towards internal bodily signals. To quantify the relative balance between external and internal processing across states, we further introduce an exteroceptive-interoceptive index derived from the ratio between AEP and HEP responses.

## Results

### Visual inspection of auditory and cardiac responses across vigilance states

We examined neural responses to external auditory stimuli (AEPs, Fig. 1A) during an isochronous sequence and to internal cardiac signals heartbeats (HEPs, Fig. 1B) during a baseline condition without auditory stimulation across wakefulness, tonic REM sleep, and phasic REM sleep. In addition to the AEPs and HEPs, neural responses to auditory and cardiac signals were quantified using the global field power (GFP), derived from evoked activity. The GFP provides an index of the strength of the evoked response irrespective of the topographic distribution of the voltage measurements across the electrode montage^30^.

**Figure 1.**
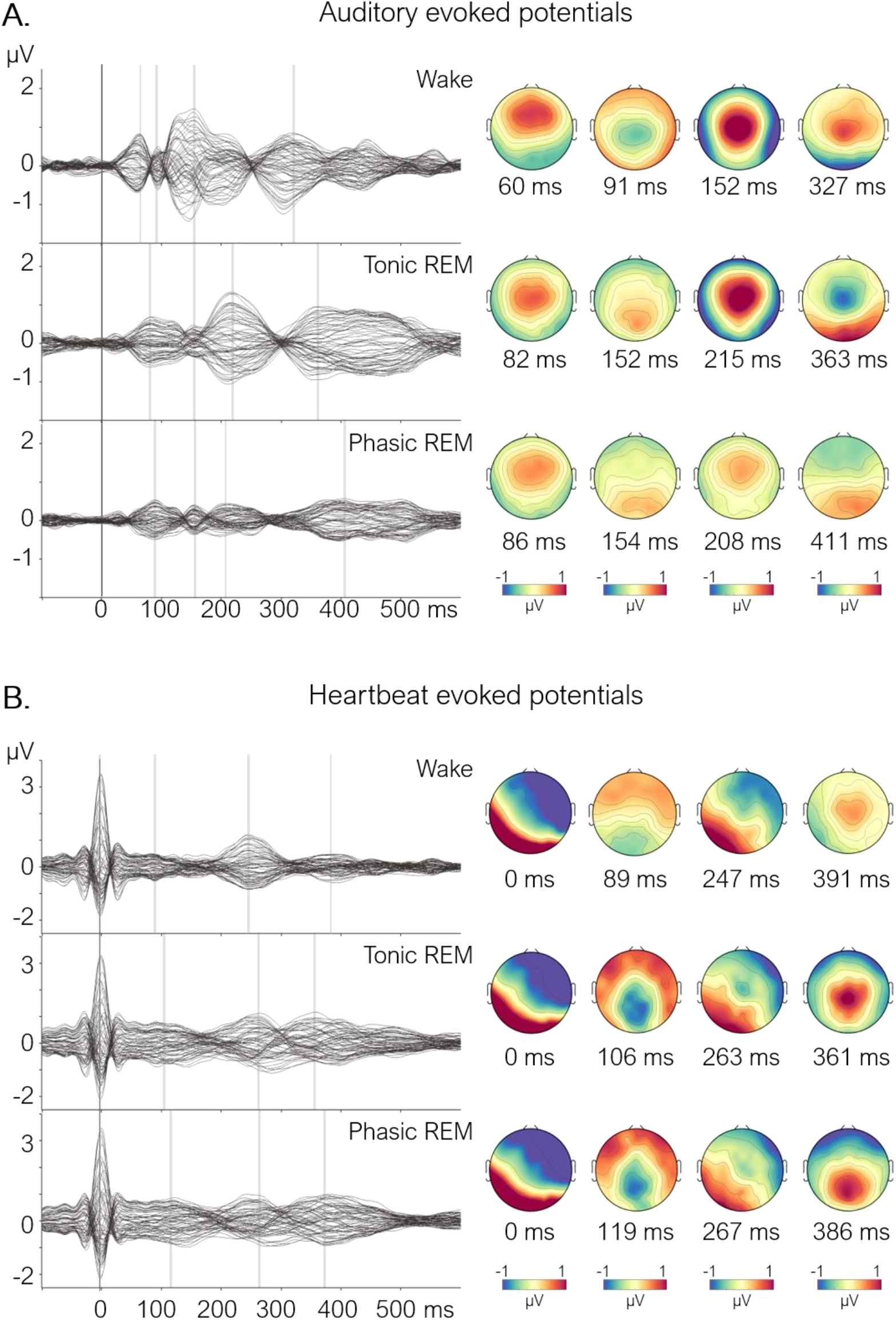
Auditory and heartbeat evoked potentials across vigilance states. **A**. Grand-average auditory evoked potentials (AEPs; n=25) shown for 62 electrodes (mastoids excluded for visualization), computed after matching the number of trials across vigilance states within each participant for wakefulness, tonic REM sleep, and phasic REM sleep. **B**. Grand-average heartbeat-evoked potentials (HEPs; n=25) obtained using the same procedure and participants. In both panels, the x-axis shows time (ms), with time 0 corresponding to sound onset (A) and to the ECG R-peak in (B); the y-axis shows amplitude (μV). Grey vertical lines indicate the latencies displayed in the scalp topographies shown on the right side of the panel.

Inspection of the grand-average AEPs revealed pronounced state-dependent changes in waveform morphology (Fig. 1A). During wakefulness, sounds elicited an early positive deflection (P50) peaking at ∼60ms over frontocentral electrodes, followed by a negative component (N100) peaking at ∼90ms maximal over centro-posterior regions. A positive component (P200) emerged at ∼150ms over central electrodes, followed by a late frontal negativity at ∼300ms.

During tonic REM sleep, the early positive component peaked later (∼80ms) and showed a centrofrontal maximum. The N100 component observed in wakefulness appeared with reversed polarity, manifesting as a positive deflection at ∼150ms. The P200 occurred later (∼215ms), followed by a late component at ∼360ms with a fronto-posterior dipolar pattern (frontal negativity and posterior positivity).

In phasic REM sleep, the temporal sequence of components resembled tonic REM sleep, but amplitudes were attenuated across the entire response compared with both wakefulness and tonic REM sleep.

In contrast, HEPs showed relatively stable waveform morphology across vigilance states (Fig. 1B). In all states, heartbeats elicited a component peaking at ∼250ms with a fronto-posterior dipolar distribution (frontal negativity and posterior positivity), followed by a later positive component at ∼380ms maximal over central electrodes.

### Statistical comparison of auditory and cardiac responses across vigilance states

Pairwise cluster-permutation tests (p<0.05, two-tailed; Fig. 2, Table S1) were employed to assess differences between wakefulness, tonic and phasic REM sleep in grand-average AEPs and HEPs and their corresponding GFP. Comparing evoked responses across vigilance states investigated differences in the magnitude and topographic distribution of the voltage measurements whereas the GFP comparison explored differences in the strength of the evoked activity^30^. Effect sizes were estimated with Cohen’s d. Control analyses using intra-sleep wakefulness i.e., periods of wakefulness during sleep, yielded comparable results, indicating that findings were not driven by differences in auditory stimulus intensity or body position between wakefulness and sleep sessions (Fig. S1).

**Figure 2.**
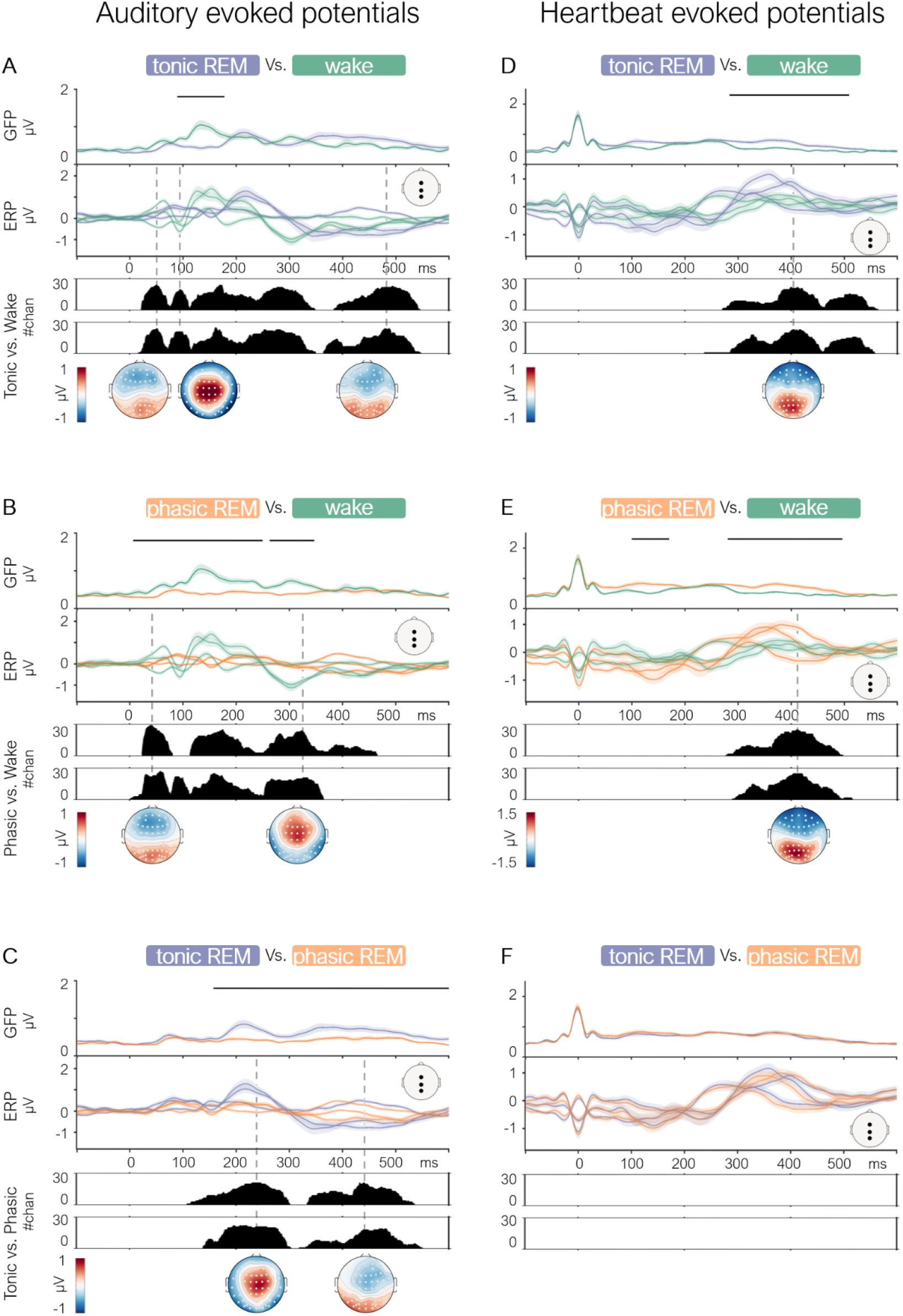
Pairwise comparison of auditory and heartbeat evoked potentials across vigilance states. Left column (A-C): Grand-average auditory evoked potentials (AEPs; n=25) comparing tonic REM sleep vs. resting wakefulness (A), phasic REM sleep vs. resting wakefulness (B) and tonic vs. phasic REM sleep (C). For each comparison, panels show (top to bottom): global field power (GFP), with black lines indicating significant time points identified by cluster permutation tests (p<0.05, two-tailed); grand-average AEPs at representative electrodes (Fz, Cz, Pz; locations shown in the scalp schematic), with green lines for wakefulness, purple for tonic REM sleep, and orange for phasic REM sleep; the number of significant electrodes over time from the cluster-based permutation analysis (upper panel: negative clusters; lower panel: positive clusters). The y-axis represents time (ms), with time 0 corresponding to sound onset. Dashed lines indicate the maximum number of electrodes within each significant cluster. Corresponding scalp topographies of AEP differences are shown below, with white dots indicating electrodes with significant results. Right column (D, E, F): Same layout and analyses for heartbeat evoked potentials (HEPs), where time 0 corresponds to ECG R peak.

**Figure 3.**
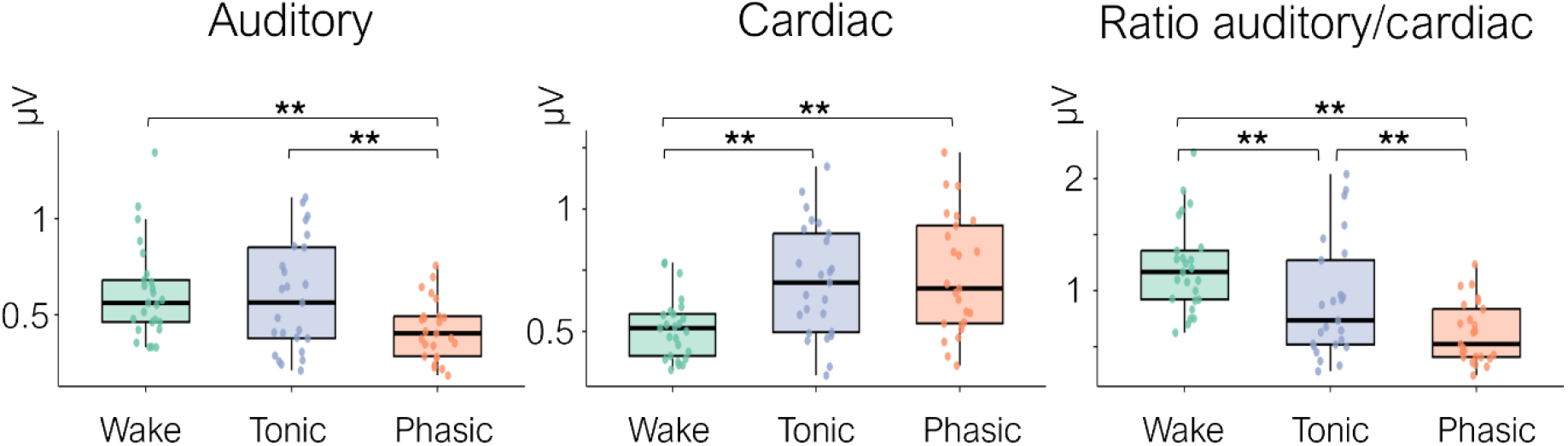
Auditory and heartbeat evoked activity and their ratio across vigilance states. Boxplots show global field power (GFP) values averaged across significant time windows for auditory-evoked potentials (AEPs; left), heartbeat-evoked potentials (HEPs; middle), and their ratio (AEP/HEP; right) across wakefulness (Wake, green), tonic REM sleep (Tonic, purple), and phasic REM sleep (Phasic, orange). Box boundaries represent the first and third quartiles, horizontal lines indicate medians, and individual points show subject-level averages. Linear mixed models were fitted for each measure with vigilance state as the fixed effect and participant as a random effect. Post-hoc pairwise comparisons were performed using Tukey adjustment. **p < 0.01

### Auditory responses

Overall, AEPs (and the corresponding GFP) showed a graded reduction across vigilance states, with the strongest responses during wakefulness, intermediate responses during tonic REM sleep, and the greatest attenuation during phasic REM sleep.

#### Tonic REM vs. Wakefulness (Fig. 2A)

GFP comparison revealed a significant negative cluster between 88 and 177 ms (p<0.01), indicating reduced response strength during tonic REM. Cluster-based analysis of AEPs identified two positive clusters (16–341ms, p<0.01, d=0.91 and 355–534ms, p<0.01, d=0.76), and three negative clusters (21–117ms, p=0.03, d=0.98; 111–340ms, p<0.01, d=0.89 and 374–532ms, p<0.01, d=0.78). Early negative clusters aligned temporally with the N100 component in wakefulness and showed a central scalp distribution. Positive clusters were more laterally distributed, while later negative clusters displayed posterior maxima with corresponding frontal positivity. These results indicate altered timing and reduced amplitude of auditory responses during tonic REM relative to wakefulness.

#### Phasic REM vs. Wakefulness AEPs (Fig. 2B)

GFP comparison revealed two significant negative clusters (5–248ms, p<0.01 and 262–346ms, p<0.01), reflecting weaker auditory responses during phasic REM. Cluster-based analyses identified two positive clusters (0–234ms, p<0.01, d=1.38 and 226–355ms, p<0.01, d=1.20) and two negative clusters (23–79ms, p=0.01, d=1.07 and 110–454ms, p<0.01, d=1.14). Spatial distribution resembled those observed in the tonic REM–wakefulness contrast, but were less pronounced, indicating stronger suppression of auditory responsiveness during phasic REM.

#### Tonic vs. Phasic REM AEPs (Fig. 2C)

GFP comparison revealed a significant positive cluster from 156 to 600 ms (p<0.01), indicating stronger auditory responses during tonic REM sleep. Cluster-based analyses identified two positive clusters (134–296ms, p<0.01, d=1.11; 311–537ms, p=0.01, d=0.81) and two negative clusters (105–294ms, p<0.01, d=1.03 and 325–522ms, p<0.01, d=0.84). These findings confirm higher auditory responsiveness during tonic compared to phasic REM sleep.

### Cardiac responses

In contrast to auditory responses, HEPs were enhanced during REM sleep relative to wakefulness.

#### Tonic REM vs. Wakefulness HEPs (Fig. 2D)

GFP analysis identified a significant positive cluster from 282 to 508 ms (p<0.01). Cluster-based comparisons of HEPs revealed a positive cluster (230–544ms, p<0.01, d=1.06) over centrofrontal electrodes and a negative cluster (264–550ms, p<0.01, d=0.85) over parieto-occipital electrodes, indicating stronger neural responses to cardiac signals during tonic REM compared to wakefulness.

#### Phasic REM vs. Wakefulness HEPs (Fig. 2E)

Two positive GFP clusters were identified (99–169ms, p<0.01; 279–495ms, p=0.01). Cluster-based analyses of HEPs revealed a positive cluster (282–503ms, p<0.01, d=1.18) and a negative cluster (270–485ms, p<0.01, d=1.10), again indicating enhanced cardiac responses during phasic REM compared to wake.

#### Tonic vs. Phasic REM HEPs (Fig. 2F)

No significant differences were observed between REM microstates in either GFP or spatiotemporal HEP distributions (p>0.05, two-tailed), indicating comparable neural processing of cardiac signals during tonic and phasic REM sleep.

### Exteroceptive-interoceptive index across vigilance states

To quantify the relative balance between external and internal sensory processing, we computed an exteroceptive–interoceptive index, defined as the ratio of AEP to HEP GFP values. Linear mixed models tested whether GFP for AEPs, HEPs, or their ratio best discriminated between vigilance states. GFP values were averaged over the longest significant time windows identified by the cluster permutation analysis (Fig. 2; AEP: ∼0–550ms, HEP: ∼100–520ms). All models included vigilance state as a fixed effect and participant as a random effect. Analyses were repeated using intra-sleep wakefulness, yielding comparable results (Fig. S2).

The exteroceptive-interoceptive index showed the strongest discriminative power across vigilance states (R^2^marginal=0.26) compared to AEPs (R^2^marginal=0.12) or HEPs (R^2^marginal=0.18) alone. Post-hoc pairwise comparisons confirmed that the index significantly differentiated all three vigilance states: wakefulness vs. tonic REM (estimate=0.30, *p*<0.01), wakefulness vs. phasic REM (estimate=0.59, *p*<0.01), and tonic vs. phasic REM (estimate=0.29, *p*<0.01). In contrast, AEPs alone failed to distinguish wakefulness from tonic REM (*p*=0.89), although they separated phasic REM from both wakefulness (estimate=0.19, *p*<0.01) and tonic REM (estimate=0.17, *p*<0.01). Similarly, HEP alone could not differentiate tonic from phasic REM (*p*=0.79), despite distinguishing wakefulness from both REM microstates (wakefulness vs. tonic: estimate=-0.19, *p*<0.01; wakefulness vs. phasic: estimate=-0.22, *p*<0.01).

Together, these results reveal a progressive shift from externally dominated sensory processing during wakefulness to internally dominated processing during phasic REM sleep.

## Discussion

The present study shows that reduced behavioural responsiveness during REM sleep does not reflect a global suppression of sensory processing, but rather a state-dependent shift in the balance between exteroceptive and interoceptive processing. Neural responses to auditory stimuli were progressively attenuated from wakefulness to tonic and phasic REM sleep. Responses to cardiac signals were preserved and even enhanced in both tonic and phasic REM microstates compared to wakefulness. Consistent with this dissociation, an exteroceptive–interoceptive index based on the ratio between auditory- and heartbeat-evoked responses reliably discriminated the three vigilance states.

Auditory responses exhibited a graded reduction across vigilance states. During tonic REM sleep, AEPs retained a waveform structure similar to wakefulness but showed delayed latencies and reduced GFP. In contrast, phasic REM sleep was characterized by markedly attenuated AEPs over extended time windows relative to both wakefulness and tonic REM sleep. These findings align with behavioural and electrophysiological evidence showing reduced responsiveness to external stimuli during phasic compared to tonic REM sleep^14,15,20,25,27^.

In contrast to auditory responses, cardiac responses showed stable morphology across vigilance states but increased amplitude during REM^24^, indicating preserved or enhanced neural processing of internal bodily signals when external responsiveness is reduced. The preservation of cardiac responses during REM sleep may also have functional implications. One possibility is that continued cortical monitoring of physiological signals supports the detection of relevant bodily changes during sleep. The brain may remain capable of detecting physiologically significant events, such as cardiovascular irregularities, that require arousal or compensatory responses. At the same time, the partial preservation of external responsiveness during tonic REM may allow intermittent monitoring of the environment.

Cardiac responses had been previously investigated in REM microstates^24,26^, although never in concomitance with auditory responses in the same participants. The differences in HEPs between wakefulness and REM sleep outlined herein are consistent with previous work^26^ however, they occurred over a shorter time window (∼350-470ms) compared to the present study (∼230-550ms). Yet, the tonic and phasic REM HEPs comparison yielded distinct findings to previous literature^26^. Simor and colleagues showed differences when comparing HEPs between the two REM microstates at ∼550-650ms while we observed no differences in the HEPs between the two REM microstates, where we focused our analysis to a post-stimulus epoch length of 500ms length.

To directly quantify the balance between external and internal sensory processing, we introduced a novel exteroceptive–interoceptive index defined as the ratio between auditory and cardiac evoked neural responses. This index accounted for more variance across vigilance states than either auditory or cardiac measures alone and successfully differentiated wakefulness, tonic, and phasic REM sleep. This index revealed a continuous shift from externally dominated processing during wakefulness toward internally dominated processing during phasic REM sleep, with tonic REM occupying an intermediate position. By jointly capturing exteroceptive and interoceptive responses, this metric provides a physiology-based measure of responsiveness that does not rely on overt behaviour.

These findings refine conventional conceptualizations of responsiveness during sleep, which have been primarily based on reactions to external stimuli. Increased arousal thresholds and reduced behavioural responses to external inputs during REM sleep have traditionally been interpreted as reflecting a generalized decrease in sensory responsiveness^14,15,20,25,27,28^. While our results confirm a reduction in neural responses to external auditory stimulation, they further show that internal bodily signals remain strongly represented in neural activity during REM sleep.

This dissociation between exteroceptive and interoceptive stimulus processing may also relate to differences in dream phenomenology across REM microstates. Tonic REM has been associated with more thought-like and less vivid mental experiences, whereas phasic REM is characterized by immersive, perceptually rich, and emotionally intense dreams^18,31^. The combination of reduced external responsiveness and enhanced internal processing during phasic REM sleep may therefore create conditions that favour internally generated experiences dominated by bodily sensations and emotions. Future work combining evoked responses with detailed dream reports could clarify whether this pattern of sensory processing directly relates to specific features of dream content^32^.

In this study, we attempted to control for potential confounding factors. Differences between wakefulness and sleep sessions, including body position for HEPs and sound intensity for AEPs, could potentially influence the observed effects. However, control analyses using intra-sleep wakefulness^26^ (Fig. S1-2) largely confirmed the results obtained during the wakefulness session, suggesting that the observed AEPs and HEPs modulations were not driven by these factors. Several reasons prompted us to utilize the separate wakefulness session. First, intra-sleep wakefulness represents a transitional state that may retain residual sleep-like features. Previous studies showed that the first minutes after awakening are associated with increased low-frequency EEG power and reduced beta activity relative to pre-sleep wakefulness^33,34^. Furthermore, intra-sleep wakefulness exhibits reduced cerebral blood flow velocity^35^ and decreased functional connectivity within sensorimotor networks resembling NREM sleep patterns^36^ in comparison to wakefulness. Second, intra-sleep wakefulness is more prone to movement-related artifacts in the EEG signal.

Finally, differences in cardiovascular dynamics between wakefulness and sleep could potentially influence HEP measurements. Although ECG activity differed between REM microstates and wakefulness (Fig. S3), these differences did not fully overlap with the time windows showing HEP modulation (Fig. 2DEF). Together with our dedicated artifact-removal procedures^37^, this supports the interpretation that the observed HEP differences reflect genuine processing rather than residual cardiac artifacts.

In summary, by leveraging the existing variability in sensory disconnection across wakefulness and REM microstates, we show that external and internal sensory processing are differentially modulated during sleep. Neural responses to external auditory stimuli progressively decrease from wakefulness to tonic and phasic REM sleep, whereas neural responses to internal cardiac signals are preserved and enhanced during REM sleep relative to wakefulness. The interoceptive-exteroceptive index provides a novel and reliable tool for outlining and assessing the mechanisms of neural responsiveness in states where awareness of the external environment is altered. These findings reveal a shift in sensory prioritization from the external environment towards internal bodily signals during REM sleep and provide a framework for investigating sensory processing in conditions where behavioural responsiveness is absent, such as anaesthesia, disorders of consciousness like coma, where neural responses to cardiac stimuli can be preserved, especially in favourable outcome patients^9^.

## Methods

### Ethics

The study (Project-ID: 2020-02373) was approved by the local ethics committee (CER-VD).

### Participants

Thirty-three healthy volunteers (16 female, age=23.42±3.48 years) with good sleep quality were included in this study, after eligibility was confirmed through a structured phone interview confirming the absence of hearing issues or psychiatric, neurological, respiratory, and cardiovascular disorders, including sleep apnoea. The Pittsburgh Sleep Quality Index and the Epworth Sleepiness Scale questionnaires ensured a regular sleep–wake schedule without excessive daytime sleepiness or sleep disturbances, respectively. All participants provided written informed consent and were not informed on the goal of the study until the end of the last experimental session.

Three of the 33 participants were excluded as they did not complete all sleep sessions. Five additional participants were excluded due to insufficient phasic REM sleep data. The final sample therefore consisted of 25 participants (14 female, age=23.24±3.65 years).

### Experimental procedures

Experimental procedures are described in detail elsewhere^37,38^. Briefly, following a cross-over design, participants underwent three consecutive overnight sleep sessions and a wakefulness session on a separate day, up to a week before or after the sleep sessions. The wakefulness session was performed in a dimly lit experimental booth equipped with sound and electrical noise attenuation. The three sleep sessions were conducted on consecutive nights in a bedroom with sound and electrical insulation. A sleep and dream diary for a week before and actigraphy monitoring (Actigraph GT3X+, ActiGraph, FL, USA) for three nights prior to the first sleep session confirmed a high sleep quality prior to participation. The first sleep session was an adaptation night, conducted for participants to familiarize themselves with sleeping at the laboratory and for the experimenters to confirm a high sleep efficiency. An auditory stimulation experiment was conducted on the second and third sleep sessions; these experimental nights are included in the study described herein. During the wakefulness session and two experimental nights, EEG signals were recorded using the 64-channel ANT Neuro EEG system (eego mylab, ANT Neuro, Hengelo, Netherlands) sampled at 1024 Hz and lab streaming layer (LSL, https://github.com/sccn/labstreaminglayer), with electrode CPz serving as the reference and AFz as the ground. ECG, vertical and horizontal EOG and submental EMG were additionally acquired.

Auditory stimulation was administered using PsychoPy and custom-made Python scripts. Three auditory regularity sequences (300 sounds each, described in detail elsewhere) and a baseline condition without auditory stimulation were administered in experimental blocks of four conditions presented in a pseudo-random order. All conditions within a block were separated by an interval of 30 seconds and the experimental block of four conditions lasted approximately 20 minutes. A pause occurred after each block of four conditions in wakefulness until a total of six blocks was administered while, in sleep, paradigm administration ensued without pauses until participants woke up the next morning. In the study described herein, we only analysed the data from the baseline to generate HEPs and the data from an isochronous sequence with fixed sound-to-sound intervals to generate AEPs. Sounds were 1000 Hz sinusoidal 16-bit stereo tones of 100 ms duration and 0 μs inter-aural time difference sampled at 44.1 KHz. A tukey window was applied to each tone. Sound stimuli were presented binaurally through in-earphones with an intensity adjusted at the single-subject level to a comfortable level, determined prior to the experimental paradigm administration. Sound intensity was different between wakefulness and the two experimental sleep sessions.

### Sleep scoring

The two experimental nights were scored by an expert sleep technician (SOMNOX SRL, Belgium) in accordance with the AASM guidelines. Tonic and phasic REM sleep were independently scored in periods of REM sleep by a second sleep expert (JC) through visual inspection, based on the occurrence of eye movement bursts. Five-second segments were labelled as phasic if the EOG signal exhibited at least two consecutive rapid eye movements (each under 500 ms in duration). Segments without any rapid eye movements were classified as tonic. Epochs containing a single rapid eye movement were considered mixed and were excluded from further analysis. Tonic REM sleep constituted a significantly larger proportion of REM sleep (mean±std: 63.67%±10.19%; range: 47.74%-87.46%) than phasic REM sleep (mean±std: 27.03%±9.03%; range: 7.61%-43.56%).

### Data analysis

Data analysis was performed in MATLAB (R2019b, The MathWorks, Natick, MA) using FieldTrip, EEGLAB, and custom scripts. Statistical analyses involving linear mixed models were performed in R Studio (4.5.1, R Core Team, 2025).

The analysis described herein was performed separately for the wakefulness session and each of the sleep sessions only for periods scored as tonic and phasic REM, as well as periods scored as intra-sleep wakefulness (Fig. S1-2). Intra-sleep wakefulness refers to epochs scored as wakefulness occurring after sleep onset (wake after sleep onset, WASO). Detailed data analysis procedures can be found in ^37^. In brief, continuous wakefulness and sleep EEG and ECG data were bandpass filtered between 0.5 and 30 Hz. ECG R-peaks were identified using the Pan-Tompkins algorithm. Artifact electrodes were excluded following a semi-automated approach implemented in FieldTrip and visual inspection. Independent component analysis (ICA) was performed to identify and remove the cardiac field artifact, as well as ocular, muscle, and sweating-related activity.

Next, cardiac-related pulse artifacts, known to have a large impact on the HEP, were detected using a quantitative analysis utilizing pairwise phase consistency (PPC) at 0-25 Hz between each ICA component and the ECG^39^. The PPC between each independent component and the ECG signal was calculated in separate time-windows of 1000 non-overlapping epochs extracted from −100 to 500 ms time-locked to the R peaks. The minimum of the median PPC values for cardiac-field artifact independent components across time-windows, participants and within each vigilance state was computed and served as the threshold for the identification of further cardiac-related components. Independent components exceeding this threshold in at least one time-window were visually inspected to confirm the presence of cardiac-related artifacts. Cardiac-related artifacts independent components as well as previously identified cardiac field artifact, ocular, muscle, and sweating components were removed, and the EEG signals were reconstructed. We also performed upcoming analyses in reconstructed data following the removal of only the cardiac field artifact, ocular, muscle, and sweating components, which produced comparable results in terms of effect sizes, temporal profiles, and scalp topographies to those described herein.

Artifact electrodes (wake: 7.4±2.4, tonic and phasic REM: 11.5±2.8) were interpolated using the spline method. Next, EEG data were segmented into non-overlapping trials between -100 and 600 ms around the sound onset in the isochronous condition to generate AEPs and between -100 and 600 ms around the R peak in the baseline condition to generate HEPs. A semi-automated trial rejection procedure using signal amplitude, standard deviation, kurtosis, z-score, and variance was performed separately for HEPs and AEPs in FieldTrip. Common average re-referencing of the EEG was performed.

AEPs and HEPs from the two experimental nights were combined (since no differences in sleep macrostructure were observed between the two nights^37^, and participants with fewer than 100 trials in wakefulness or tonic REM sleep, phasic REM sleep, or intra-sleep wakefulness were excluded from further analyses. Three participants were excluded from the intra-sleep wakefulness analysis due to insufficient trial counts or excessive movement artifacts. Trials were matched across vigilance states, separately for AEPs and HEPs within each participant (mean±std trial numbers: AEPs=502.44±274.90; HEPs=558.48±292.81). Grand-average AEPs and HEPs were computed for each participant and vigilance state, and GFP was calculated from these averages as the standard deviation of the evoked activity across channels at each time point.

### Statistical analysis

Non-parametric cluster-based permutation statistics (p<0.05, two-tailed; 5000 permutations) contrasted the AEPs, HEPs and their GFPs for pairs of the vigilance states of interest (wake, tonic REM sleep, phasic REM sleep, intra-sleep wakefulness) over the entire trial length (-100 to 600 ms). As a control, the same analysis was performed for the ECG to ensure that observed differences between vigilance states were not driven by differences in the ECG signal. The Cohen’s d statistic at the latencies where the largest number of electrodes showed significant differences following permutation testing was used to calculate effect sizes.

Linear mixed models were fitted to compare the discriminative ability of AEP, HEP, and their ratio (exteroceptive-interoceptive index) across vigilance states (wakefulness, tonic REM sleep, phasic REM sleep). GFP values were averaged over the longest significant time windows from cluster permutation analyses and used as dependent variables. All models included vigilance state as a fixed effect and subject as a random intercept and were fitted using the lme4 package in R. Model comparisons were based on marginal R^2^ (variance explained by vigilance state), calculated with R’s *performance* package. Post-hoc pairwise comparisons used Tukey-adjusted estimated marginal means (R’s *emmeans* package).

## Data and code availability

Data and analysis code are available by the corresponding author upon reasonable request. The code for the experimental paradigm administration (https://github.com/fcbg-platforms/eeg-cardio-audio-sleep/tree/maint/0.3) is publicly available.

## Acknowledgements

The authors thank Matthieu Scheltienne from the Campus Biotech Foundation in Geneva for designing the code for the experimental paradigm administration, and Gwenael Birot and Virginie Sterpenich for useful feedback and guidance with regards to experimental procedures. Finally, we are grateful to Alice Clerget, Erin Mahan and Marie Zocca for assistance with participant recruitment and data acquisition.

This work was supported by the Bertarelli Foundation grant ‘Catalyst’ to MDL and SS, the Swiss National Science Foundation (grant 32003B_212981), the Eurostars project E!3489 to MDL, and the University of Lausanne ‘Transition Grant’ to AP.

## Author contributions

**Jacinthe Cataldi:** Data curation, Methodology, Formal analysis, Writing - original draft; **Andria Pelentritou:** Conceptualization, Data acquisition, Data curation, Methodology, Formal analysis, Writing - original draft; **Sophie Schwartz:** Conceptualization, Resources, Funding acquisition, Project administration, Writing - review and editing; **Marzia De Lucia:** Conceptualization, Methodology, Resources, Funding acquisition, Project administration, Supervision, Writing - original draft.

## Declaration of interests

The authors declare no competing interests.

## Notes

### Competing Interest Statement

The authors have declared no competing interest.

