## Supplementary Materials for "Sensory processing reallocation from external to internal signals in REM sleep"

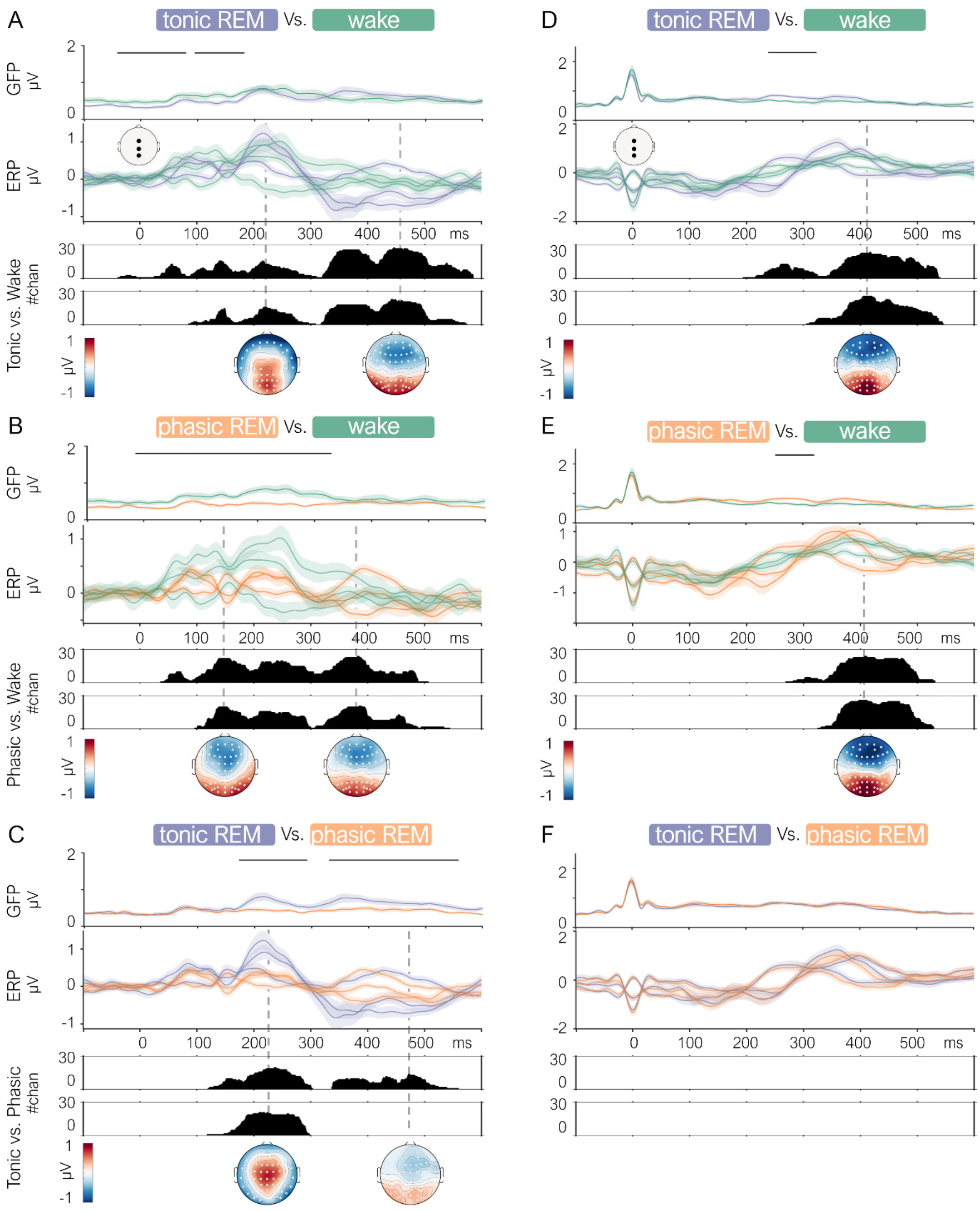


**Figure S1. Comparison of auditory and heartbeat evoked potentials across vigilance states.** Left column (A, B, C): Grand-average (n=22) auditory evoked potentials (AEPs) pair-wise comparison of tonic REM sleep vs. infra-sleep wakefulness (A), phasic REM sleep vs. infra-sleep wakefulness (B) and tonic vs. phasic REM sleep (C). For each comparison, the panels show (top to bottom): grand-average GFP with the black line indicating the significant timepoints after cluster permutation statistical analysis ($p<0.05$, two-tailed); grand-average AEP, displayed for three representative electrodes (Fz, Cz, Pz; locations shown in white topographic schema; green lines for wakefulness, purple for tonic and orange for phasic REM sleep); cluster permutation statistical analysis results contrasting AEPs between vigilance states shown as the number of significant electrodes ($p<0.05$, two-tailed), where the upper plot displays negative clusters and the lower plot displays positive clusters of electrodes. The y-axis represents time (ms), with zero corresponding to sound onset. Dashed lines indicate the maximum number of electrodes for each significant cluster, with corresponding topographical distributions of AEPs differences between vigilance states shown at the bottom. White dots on topographic maps indicate significant electrodes. Right column (D, E, F): Same layout and analyses for heartbeat evoked potentials (HEPs), where the zero of the y-axis corresponds to R-peak latency of the ECG signal.

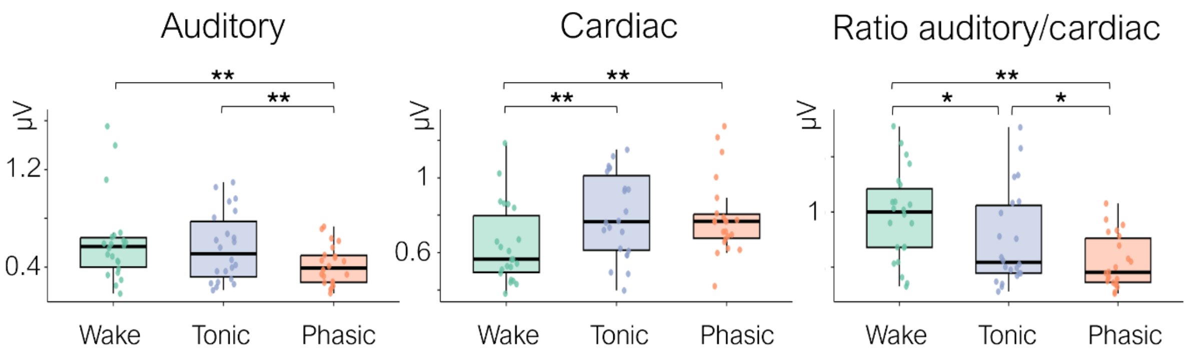


**Figure S2. Auditory and heartbeat evoked activity and their ratio across vigilance states**. Boxplots showing global field power (GFP) values averaged over significant time windows (AEP: ~ -40–570ms; HEP: 240–340ms; see Fig. S1) for auditory-evoked potentials (AEP, left), heartbeat-evoked potentials (HEP, middle), and their ratio (AEP/HEP, right) across infra-sleep wakefulness (Wake, green), tonic REM (Tonic, purple), and phasic REM (Phasic, orange). Box boundaries represent the first and third quartiles, horizontal lines indicate medians, and individual points show subject-level averages. Linear mixed models with vigilance state as fixed effect and subject as random effect were fitted for each measure. Post-hoc pairwise comparisons with Tukey adjustment are indicated above each panel. **p < 0.01 ; *p<0.05

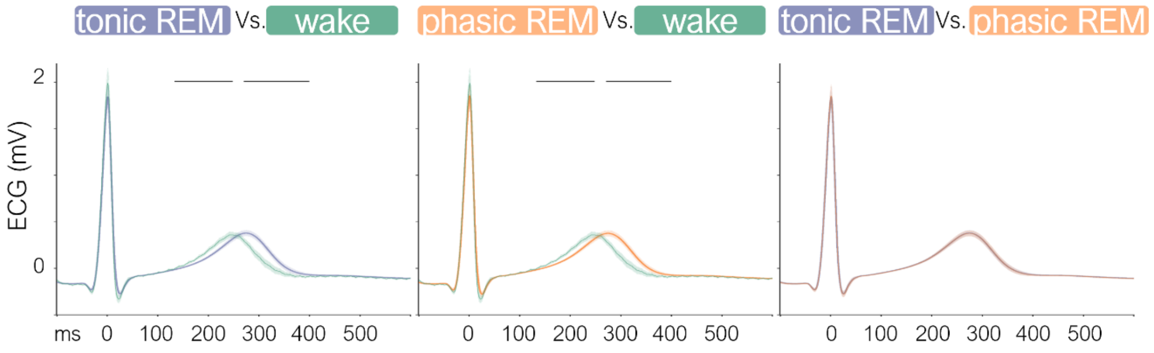


**Figure S3. ECG comparison across vigilance states. A.** Grand average ECG comparison between vigilance states (n=25): wakefulness (green), tonic (purple) and phasic (orange) REM sleep. The waveforms were obtained averaging 100 trials for each vigilance state. The black line in each contrast indicates the significant timepoints after a cluster permutation statistical analysis (p<0.05, two-tailed). The x-axis represents time, with zero corresponding to R-peak latency; the y-axis represents amplitude. ECG waveforms comparisons between vigilance states showed difference between wakefulness and the two REM microstates. Tonic REM vs. wakefulness: one negative (132-247ms, p<0.01) and one positive cluster (269-398ms, p<0.01) emerged. In phasic REM vs. wakefulness: one negative (132-247ms, p<0.01) and one positive cluster (270-399ms, p<0.01) emerged.

**Table S1. Summary of statistical results for the comparison of auditory and heartbeat evoked potentials across vigilance states.** Each row corresponds to a significant cluster ($p<0.05$, two-tailed) identified using a cluster-based permutation test. For each cluster, the table reports the contrast type (auditory evoked potentials - AEP or heartbeat evoked potentials - HEP; global field power - GFP or spatio-temporal evoked reponses comparison), the two compared conditions, the effect direction (positive or negative), the time window of significance in ms (start–end), the p-value (p), the effect size - Cohen's d (d), and the corresponding panel in Fig. 2 where the effect is illustrated.

| N | measure type | conditions | direction | start (ms) | end (ms) | p | d | Fig. 2 |
|---|---|---|---|---|---|---|---|---|
| | | | | AEP | | | | |
| 1 | GFP | tonic vs. Wake | neg | 88 | 117 | 4,4E-03 | | A |
| 2 | spatio-temporal | tonic vs. wake | pos | 16 | 341 | 2,0E-04 | 0,91 | A |
| 3 | spatio-temporal | tonic vs. wake | pos | 355 | 534 | 6,4E-03 | 0,76 | A |
| 4 | spatio-temporal | tonic vs. wake | neg | 21 | 117 | 2,5E-02 | 0,98 | A |
| 5 | spatio-temporal | tonic vs. wake | neg | 111 | 340 | 2,0E-04 | 0,89 | A |
| 6 | spatio-temporal | tonic vs. wake | neg | 374 | 532 | 4,8E-03 | 0,78 | A |
| 7 | GFP | phasic vs. wake | neg | 5 | 248 | 4,0E-04 | | B |
| 8 | GFP | phasic vs. wake | neg | 262 | 346 | 8,0E-03 | | B |
| 9 | spatio-temporal | phasic vs. wake | pos | 0 | 234 | 2,0E-04 | 1,38 | B |
| 10 | spatio-temporal | phasic vs. wake | pos | 226 | 355 | 6,0E-04 | 1,20 | B |
| 11 | spatio-temporal | phasic vs. wake | neg | 23 | 79 | 1,0E-02 | 1,07 | B |
| 12 | spatio-temporal | phasic vs. wake | neg | 110 | 454 | 2,0E-04 | 1,14 | B |
| 13 | GFP | tonic vs. phasic | pos | 156 | 600 | 4,0E-04 | | C |
| 14 | spatio-temporal | tonic vs. phasic | pos | 134 | 296 | 1,2E-03 | 1,11 | C |
| 15 | spatio-temporal | tonic vs. phasic | pos | 311 | 537 | 1,0E-02 | 0,81 | C |
| 16 | spatio-temporal | tonic vs. phasic | neg | 105 | 294 | 1,4E-03 | 1,03 | C |
| 17 | spatio-temporal | tonic vs. phasic | neg | 325 | 522 | 4,2E-03 | 0,84 | C |
| | | | | HEP | | | | |
| 18 | GFP | tonic vs. wake | pos | 282 | 508 | 8,0E-05 | | D |
| 19 | spatio-temporal | tonic vs. wake | pos | 230 | 544 | 2,0E-04 | 1,06 | D |
| 20 | spatio-temporal | tonic vs. wake | neg | 264 | 550 | 2,1E-04 | 0,85 | D |
| 21 | GFP | phasic vs. wake | pos | 99 | 169 | 4,0E-04 | | E |
| 22 | GFP | phasic vs. wake | pos | 279 | 495 | 1,0E-02 | | E |
| 23 | spatio-temporal | phasic vs. wake | pos | 282 | 503 | 2,0E-04 | 1,18 | E |
| 24 | spatio-temporal | phasic vs. wake | neg | 270 | 485 | 4,0E-04 | 1,10 | E |
